# Peroxidasin expression is increased in intratumoural capillaries and in proximal tubular cells adjacent to clear cell renal cell carcinoma

**DOI:** 10.1101/2024.11.15.623667

**Authors:** Roberto Silva, Jorge Reis Almeida, Ana Rita Coelho, Isabel Brandão, Bárbara Gomes, Inês Soares Alencastre, João Paulo Oliveira

**Affiliations:** Pathology Department of the Faculty of Medicine, University of Porto, Alameda Prof. Hernâni Monteiro, 4200-319 Porto, Portugal; ULS São João, EPE, Alameda Prof. Hernâni Monteiro, 4200-319 Porto, Portugal; i3S – Instituto de Investigação e Inovação em Saúde, University of Porto, R. Alfredo Allen 208, 4200-135 Porto, Portugal; Faculty of Medicine/, University of Porto / RISE – Laboratório Associado, Alameda Prof. Hernâni Monteiro, 4200-319 Porto, Portugal

**Keywords:** Peroxidasin, Clear cell renal cell carcinoma, Oncocytoma, Angiogenesis

## Abstract

Peroxidasin (PXDN), is a heme peroxidase with a critical role in the crosslinking of type IV collagen, being essential for basement membrane integrity. In many cancers, PXDN overexpression has been linked with processes associated with poor disease outcomes, namely the impairment of endothelial functions, pathogenesis of vascular diseases, vascular remodelling, apoptosis, and tissue fibrosis. In RCC, almost nothing is so far known. In this work we characterized the expression of PXDN in tumour and non-neoplastic adjacent tissue from clear cell renal cell carcinoma (ccRCC) and renal oncocytoma by immunohistochemistry. Results showed a significant increase of PXDN expression in intratumoural capillaries of ccRCC in comparison with oncocytoma, and an increase in the proximal tubular epithelial cells of non-neoplastic adjacent parenchyma from ccRCC cases, in comparison with the same area from oncocytomas. Overall, our results suggest, for the first time, that PXDN may have a role in tumour angiogenesis that may favour metastases and tumour malignity, being also enrolled in the crosstalk between the tumour and adjacent tissue, that deserves further investigation.

**Highlights:**

- Peroxidasin is increased in ccRCC microvasculature comparing to oncocytoma.
- Peroxidasin is increased in proximal tubules of ccRCC kidney adjacent parenchyma.
- Is Peroxidasin a modulator of tumour microenvironment promoting angiogenesis?

## Introduction

Peroxidasin (PXDN) is a haem-peroxidase involved in distinct physiological processes, including extracellular matrix (ECM) formation, stabilization of collagen IV (COL4) network in basement membranes, and host defence mechanisms against bacterial pathogens [1]. In cardiovascular diseases associated with increased oxidative stress, such as atherosclerosis, PXDN contributes to endothelial dysfunction and vascular remodelling, and it regulates fibrogenic responses in mouse models of myocardial infarction, chronic liver disease and obstructive nephropathy.

PXDN appears to play a role in the progression of numerous cancers [2], by promoting tumour angiogenesis and invasiveness; yet little is known about its involvement in renal cell carcinoma (RCC). PXDN protein expression is significantly increased in RCC, and the median *PXDN* RNA expression levels are 6- to 8-fold higher in clear cell (ccRCC) and papillary subtypes, relative to chromophobe RCC [3].

Overall, ccRCC accounts for the large majority (>70%) of all RCC cases. The cells of origin in ccRCC are proximal tubular epithelial cells (PTEC), but the molecular pathways underlying the transition to the malignant phenotype remain unclear [4]. By single-cell transcriptomics of ccRCC tissue [5], high expression of *PXDN* transcripts, along with other genetic markers of the tip cell angiogenic phenotype, was identified in a previously uncharacterized vasculature subpopulation, also enriched for epithelial-mesenchymal transition (EMT) pathway genes associated with worse patient survival; these cells further exhibited increased expression of ECM constituents, including pro-angiogenic and potentially pro-metastatic perlecan and COL4. Furthermore, *PXDN* is one of 12 highly correlated genes defining a transcriptomic ccRCC immune escape signature to treatment with immune checkpoint inhibitors [6]. Altogether, an increasing number of studies suggest that PXDN may contribute to the modulation of ccRCC tumour microenvironment and cell-cell communication warranting further research, namely addressing its value as a potential prognostic biomarker and/or therapeutic target.

The aim of this study was to compare the expression of PXDN by immunohistochemistry (IHC), in ccRCC, kidney oncocytomas, and histologically normal parenchyma adjacent to either tumour.

## Materials and Methods

### Ethical approval

This study was conducted in accordance with the principles of the Declaration of Helsinki and its protocol was reviewed and approved by the Health Ethics Commission and the Data Protection Officer at *Centro Hospitalar Universitário de São João* (CHUSJ), Porto, Portugal (ref. CE-04-2023).

### Cohort selection

Formalin-fixed paraffin-embedded surgical specimens of ccRCC (n = 12) and oncocytoma (n = 11) surrounded by at least 1 cm of non-neoplastic parenchyma, were retrieved from the archives of the CHUSJ Pathology Department. None of the cancer patients had undergone radiotherapy or chemotherapy before resection. Tumours were staged according to the TNM system [7]. Relevant demographic and clinicopathological features of the 23 patients are presented in Table 1.

**Table 1.**
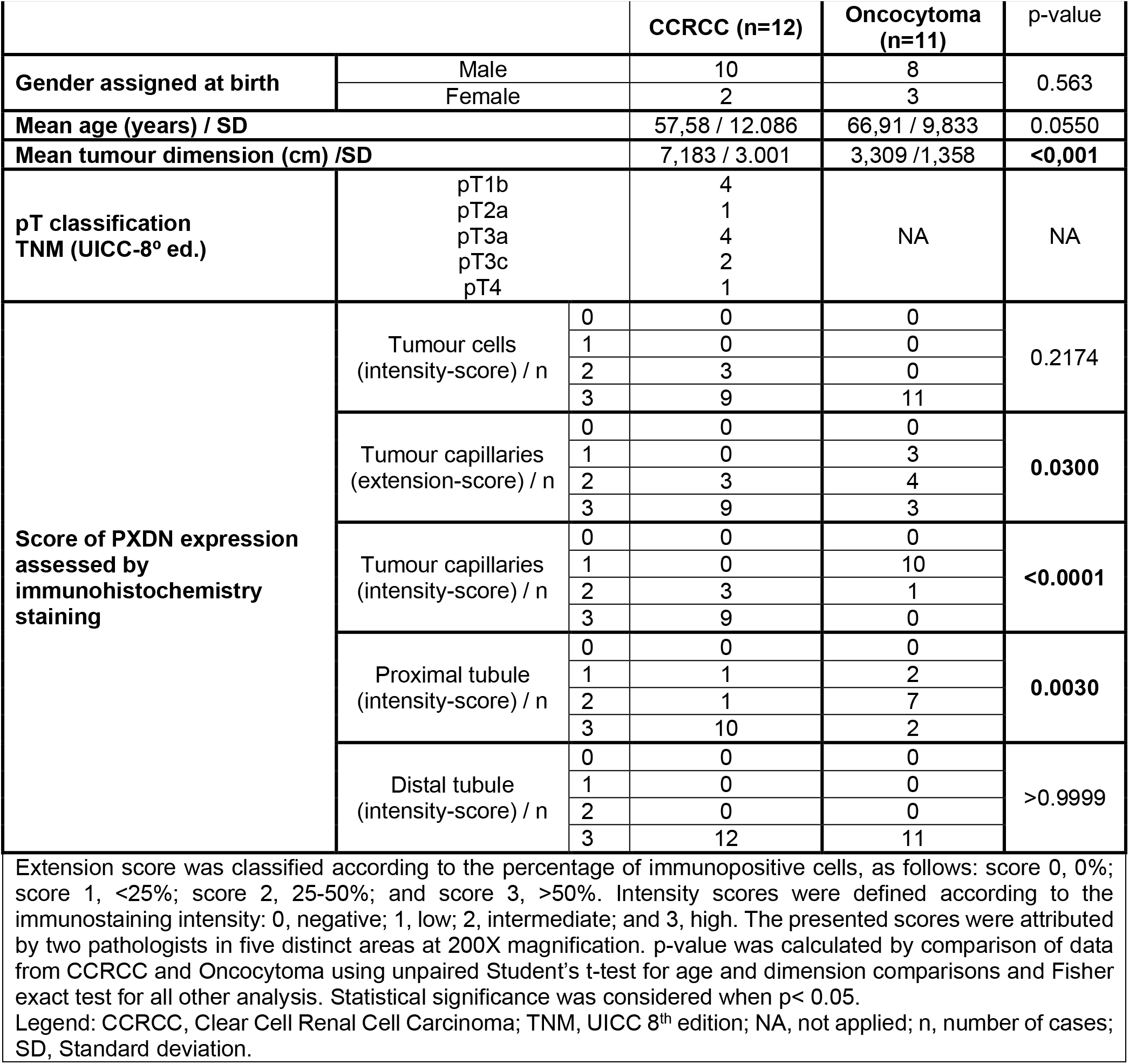
Clinicopathological features.

| Table 1. Clinicopathological features |  |  |  |  |  |
| --- | --- | --- | --- | --- | --- |
|  |  |  | CCRCC (n=12) | Oncocytoma (n=11) | p-value |
| Gender assigned at birth | Male |  | 10 | 8 | 0.563 |
|  | Female |  | 2 | 3 |  |
| Mean age (years) / SD |  |  | 57,58 / 12.086 | 66,91 / 9,833 | 0.0550 |
| Mean tumour dimension (cm) /SD |  |  | 7,183 / 3.001 | 3,309 /1,358 | <0,001 |
| pT classification<br>TNM (UICC-8<sup>o</sup> ed.) | pT1b |  | 4 | NA | NA |
|  | pT2a |  | 1 |  |  |
|  | pT3a |  | 4 |  |  |
|  | pT3c |  | 2 |  |  |
|  | pT4 |  | 1 |  |  |
| Score of PXDN expression<br>assessed by<br>immunohistochemistry<br>staining | Tumour cells<br>(intensity-score) / n | 0 | 0 | 0 | 0.2174 |
|  |  | 1 | 0 | 0 |  |
|  |  | 2 | 3 | 0 |  |
|  |  | 3 | 9 | 11 |  |
|  | Tumour capillaries<br>(extension-score) / n | 0 | 0 | 0 | 0.0300 |
|  |  | 1 | 0 | 3 |  |
|  |  | 2 | 3 | 4 |  |
|  |  | 3 | 9 | 3 |  |
|  | Tumour capillaries<br>(intensity-score) / n | 0 | 0 | 0 | <0.0001 |
|  |  | 1 | 0 | 10 |  |
|  |  | 2 | 3 | 1 |  |
|  |  | 3 | 9 | 0 |  |
|  | Proximal tubule<br>(intensity-score) / n | 0 | 0 | 0 | 0.0030 |
|  |  | 1 | 1 | 2 |  |
|  |  | 2 | 1 | 7 |  |
|  |  | 3 | 10 | 2 |  |
|  | Distal tubule<br>(intensity-score) / n | 0 | 0 | 0 | >0.9999 |
|  |  | 1 | 0 | 0 |  |
|  |  | 2 | 0 | 0 |  |
|  |  | 3 | 12 | 11 |  |
| Extension score was classified according to the percentage of immunopositive cells, as follows: score 0, 0%; score 1, <25%; score 2, 25-50%; and score 3, >50%. Intensity scores were defined according to the immunostaining intensity: 0, negative; 1, low; 2, intermediate; and 3, high. The presented scores were attributed by two pathologists in five distinct areas at 200X magnification. p-value was calculated by comparison of data from CCRCC and Oncocytoma using unpaired Student's t-test for age and dimension comparisons and Fisher exact test for all other analysis. Statistical significance was considered when p< 0.05.<br>Legend: CCRCC, Clear Cell Renal Cell Carcinoma; TNM, UICC 8<sup>th</sup> edition; NA, not applied; n, number of cases; SD, Standard deviation. |  |  |  |  |  |

### Immunohistochemistry

Three-µm serial sections, representative of tumour and non-neoplastic adjacent tissue, were processed for IHC according to a previously described protocol [8]. Briefly, the slides were deparaffinised and rehydrated, and antigen retrieval was carried out with 10 mM citrate buffer, pH 6.0, at 98ºC for 20 min; after permeabilization with 0.1% Triton X-100 and blocking with 10% normal horse serum, the slides were incubated with anti-PXDN antibody (1:100) (Abbexa; Cambridge, UK), overnight at 4ºC; finally, following quenching of endogenous peroxidase activity, the slides were incubated with the Lab Vision UltraVision Large Volume Detection System: anti-Polyvalent, HRP (Thermo Fisher Scientific Anatomical Pathology, Cheshire, UK) IHC staining kit, using diaminobenzidine (Dako, Carpinteria, CA, USA) as the peroxidase chromogen, and counterstained with hematoxylin (Leica Biosystems, Richmond, USA). Images were acquired in an Olympus BX43 microscope (Japan), using the Olympus CellSens Entry 1.12 software platform. In negative controls, the primary antibodies were omitted and replaced by the antibody dilution reagent (data not shown).

Extent and intensity of PXDN expression were semiquantitatively assessed in neoplastic cells and adjacent non-neoplastic tubular epithelial cells, as well as in endothelial cells of intratumoural capillaries. The extent of immunostaining was scored according to the percentage of stained cells, as follows: score 0, 0%; score 1, <25%; score 2, 25-50%; and score 3, >50%; the intensity score was rated from 0 to 3, respectively for “negative”, “low”, “intermediate” and “high” expression. Each specimen was scored by two pathologists (A.C. and R.S.) in five distinct areas (*hot spots)* at 200x magnification, and the resultant five scores were averaged and rounded up to the nearest integer.

### Statistical analysis

Statistical analyses were performed by IBM SPSS statistics version 29.0 (IBM SPSS, Chicago, IL, USA). The unpaired Student’s t-test was used to compare the patient’ ages and the tumour dimensions, and the Fisher exact test for all other comparisons; p-values <0.05 were considered statistically significant.

## Results

The ccRCC specimens were from ten males and two females, with median age of 61 years; all tumours were unifocal, with mean largest diameter of 7.2 cm; pathological stage was pT1b in 4 cases, pT2a in 1 case, pT3a in 4 cases, pT3c in 2 cases and pT4 in 1 case. The oncocytoma specimens were from eight males and three females, with a median age of 70 years; the mean of the largest tumour diameter was 3.3 cm, and two patients had multifocal tumours. None of the ccRCC patients had metastatic disease, and no tumour-related deaths were reported over a median follow-up of two years. In both the ccRCC and oncocytoma specimens, >50% of the tumour cells exhibited PXDN immunostaining, but the intratumoural capillaries of the former showed significantly higher PXDN staining extent and intensity (p=0.030 and p<0.001, respectively) (Figure_1 A1 and B1, Table1). In the proximal and distal tubular epithelium of tumour-adjacent histologically normal parenchyma, the extent of PXDN expression scored “3” in both the ccRCC and oncocytoma specimens. However, PXDN expression intensity in PTEC of the non-neoplastic parenchyma adjacent to ccRCC was significantly higher than in PTEC of normal parenchyma adjacent to oncocytoma (p=0.030) (Figure 1 A2 and B2, Table1); contrastingly, the intensity of PXDN expression in the distal tubular epithelium scored 3 (Table 1) in both cases. No statistical relationship was identified between PXDN expression and the ccRCC TNM stage.

**Figure 1.**
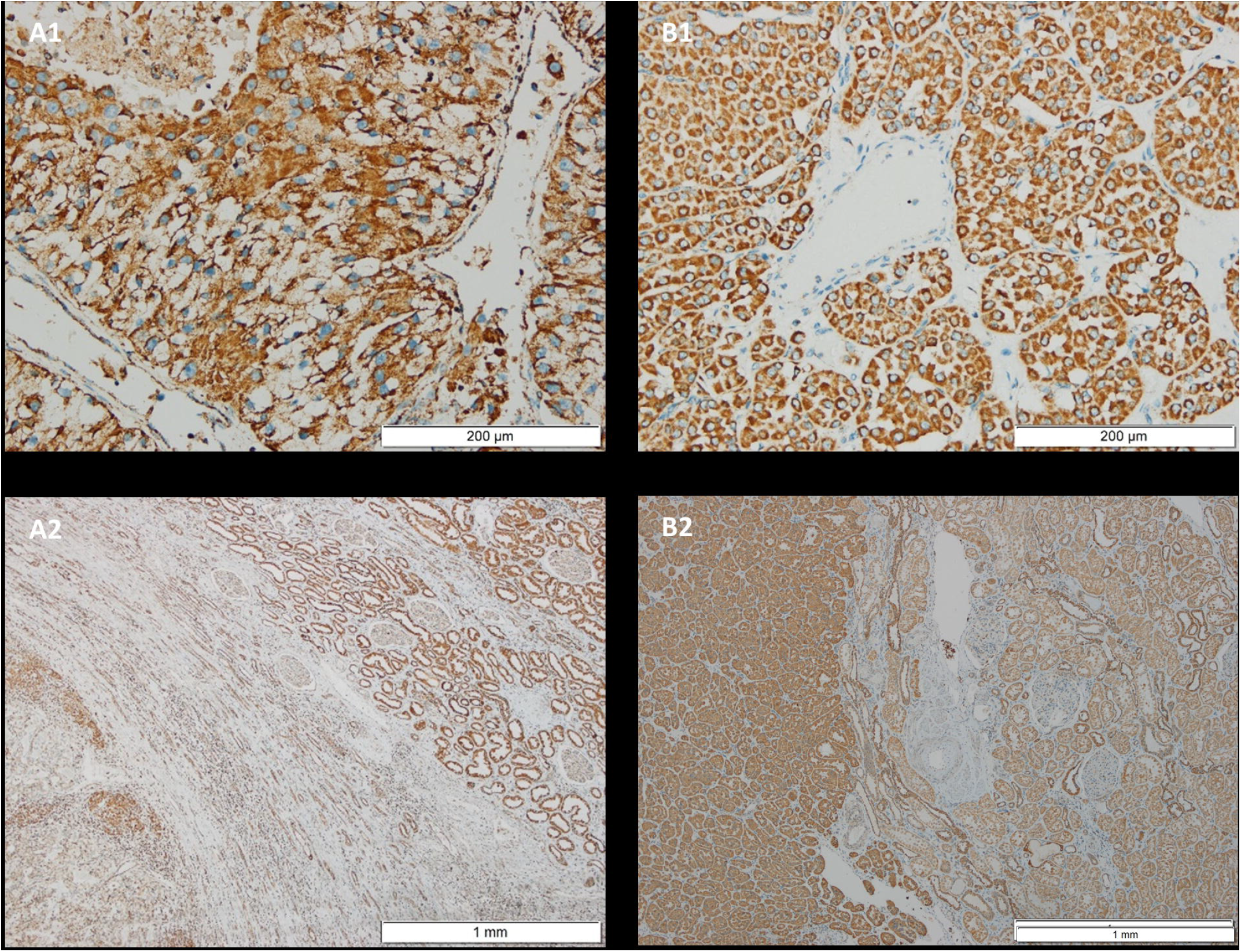
Micrographs of PXDN staining in CCRCC and oncocytoma tumour tissue and adjacent parenchyma (>1cm). Details of the histology and staining methods are provided in Materials and Methods. Scale bars: [A1, B1] 200x, [A2, B2] 40x.

## Discussion

Results of recent studies in several human cancers suggest that PXDN may have a role in the mechanisms supporting tumour growth, progression, invasion, and metastatic dissemination, including angiogenesis and EMT, and may even serve as a diagnostic biomarker for thyroid anaplastic carcinoma [2, 9-12].

In the present study, we aimed to characterize the relative PXDN expression in distinct cell-type compartments of ccRCC, as compared to a benign kidney neoplasia (oncocytoma) and non-neoplastic kidney tissue, using an IHC approach. The significantly increased microvascular PXDN expression in ccRCC as compared to oncocytoma was a most relevant finding, in line with the putative role of PXDN in tumoural angiogenesis [5], as well with experimental data from single-cell transcriptome analysis of lung tumour endothelial cells [13]. Although PXDN expression in tumoural capillaries has been previously described in ccRCC and in brain tumours [14, 15] to the best of our knowledge this is the first evidence of higher PXDN protein expression in the microvasculature of a malignant compared to a non-malignant kidney neoplasm. Considering the critical role of PXDN in the cross-linking of COL4, which is a major component of ECM and basement membranes, associated with pro-angiogenic and pro-metastatic phenotypes in several tumours [9], and the pro-angiogenic peroxidase activity of PXDN [14], our data support the hypothesis that PXDN may indeed be part of the tumour growth promoting factors in the ccRCC tumour microenvironment. An additional provocative finding, deserving further research, was the significantly higher intensity of PXDN immunostaining of the PTEC in the non-neoplastic parenchyma adjacent to ccRCC, as compared to the normal parenchyma adjacent to oncocytoma, suggesting the existence of a particular interface between the ccRCC microenvironment and the adjacent proximal tubules. Finally, the data on PXDN expression in the tumoural cells of kidney oncocytoma are also novel.

In conclusion, taking into consideration our results and the previously available data, we postulate that PXDN might be a modulator of the tumour microenvironment, influencing ECM characteristics (like integrity and stiffness) and angiogenesis, which favour tumour growth and metastasis. The confirmation of this hypothesis warrants additional research. Exploring the functional role and potential pathophysiological significance of PXDN in ccRCC could pave the way for new research in kidney cancer, potentially identifying a new biomarker or therapeutic target.

## Glossary

ccRCC: Clear cell renal cell carcinoma
COL4: Type 4 collagen
ECM: Extracellular matrix
EMT: Epithelial-mesenchymal transition
PTEC: Proximal tubular epithelial cells
PXDN: Peroxidasin
RCC: Renal cell carcinoma
TNM: Tumour nodes metastasis

## Author contributions

R.S, J.R.A, I.S.A. and J.P.O. conceived, designed, and supervised the research project; A.R.C., J.R.A. and R.S. collected and validated the kidney tissue specimens for study; B.G. and I.B. provided histopathology technical support; J.R.A. and R.S. analysed the slides; A.R.C. and R.S. did the statistical analyses; I.S.A, R.S., J.R.A and J.P.O analysed the data; I.S.A., R.S. and J.P.O. wrote the draft manuscript. All authors discussed the results and read and approved the final manuscript. The corresponding author had full access to the data in the study and final responsibility for the decision to submit for publication.

## Acknowledgements

The kidney tissue samples used in this study were kindly provided by the Department of Anatomic Pathology of *Centro Hospitalar Universitário de São João* (CHUSJ), Porto, Portugal

## Ethics approval

This study was conducted in accordance with the Declaration of Helsinki. The research protocol was reviewed and approved by the institutional Health Ethics Commission (reference number CE-04-2023) and Data Protection Officer of *Centro Hospitalar Universitário de São João* (CHUSJ), Porto, Portugal.

## References

[1] G. Cheng, R. Shi, Mammalian peroxidasin (PXDN): From physiology to pathology, Free Radic Biol Med, 182 (2022) 100–107.

[2] K. Wyllie, V. Panagopoulos, T.R. Cox, The role of peroxidasin in solid cancer progression, Biochem Soc Trans, 51 (2023) 1881–1895.

[3] M. Uhlen, C. Zhang, S. Lee, E. Sjostedt, L. Fagerberg, G. Bidkhori, R. Benfeitas, M. Arif, Z. Liu, F. Edfors, K. Sanli, K. von Feilitzen, P. Oksvold, E. Lundberg, S. Hober, P. Nilsson, J. Mattsson, J.M. Schwenk, H. Brunnstrom, B. Glimelius, T. Sjoblom, P.H. Edqvist, D. Djureinovic, P. Micke, C. Lindskog, A. Mardinoglu, F. Ponten, A pathology atlas of the human cancer transcriptome, Science, 357 (2017).

[4] A.M. Raghubar, M.J. Roberts, S. Wood, H.G. Healy, A.J. Kassianos, A.J. Mallett, Cellular milieu in clear cell renal cell carcinoma, Front Oncol, 12 (2022) 943583.

[5] J. Zvirblyte, J. Nainys, S. Juzenas, K. Goda, R. Kubiliute, D. Dasevicius, M. Kincius, A. Ulys, S. Jarmalaite, L. Mazutis, Single-cell transcriptional profiling of clear cell renal cell carcinoma reveals a tumor-associated endothelial tip cell phenotype, Commun Biol, 7 (2024) 780.

[6] M. Golkaram, F. Kuo, S. Gupta, M.I. Carlo, M.L. Salmans, R. Vijayaraghavan, C. Tang, V. Makarov, P. Rappold, K.A. Blum, C. Zhao, R. Mehio, S. Zhang, J. Godsey, T. Pawlowski, R.G. DiNatale, L.G.T. Morris, J. Durack, P. Russo, R.R. Kotecha, J. Coleman, Y.B. Chen, V.E. Reuter, R.J. Motzer, M.H. Voss, L. Liu, E. Reznik, T.A. Chan, A.A. Hakimi, Spatiotemporal evolution of the clear cell renal cell carcinoma microenvironment links intratumoral heterogeneity to immune escape, Genome Med, 14 (2022) 143.

[7] M.K.G. James D. Brierley, Christian Wittekind TNM Classification of Malignant Tumours, 8th Edition, Wiley2016.

[8] R.S. Isabel Brandão, Eduardo Conde, Bárbara Gomes, Paula Sampaio, Ana Costa Braga, Jorge Reis Almeida, Inês Soares Alencastre*, João Paulo Oliveira*, Characterization of peroxidasin expression in histologically normal human adult and fetal kidney tissue, bioRxiv (2023).

[9] X. Zhou, Q. Sun, C. Xu, Z. Zhou, X. Chen, X. Zhu, Z. Huang, W. Wang, Y. Shi, A systematic pan-cancer analysis of PXDN as a potential target for clinical diagnosis and treatment, Front Oncol, 12 (2022) 952849.

[10] J. Dougan, O. Hawsawi, L.J. Burton, G. Edwards, K. Jones, J. Zou, P. Nagappan, G. Wang, Q. Zhang, A. Danaher, N. Bowen, C. Hinton, V.A. Odero-Marah, Proteomics-Metabolomics Combined Approach Identifies Peroxidasin as a Protector against Metabolic and Oxidative Stress in Prostate Cancer, Int J Mol Sci, 20 (2019).

[11] Y.Z. Zheng, L. Liang, High expression of PXDN is associated with poor prognosis and promotes proliferation, invasion as well as migration in ovarian cancer, Ann Diagn Pathol, 34 (2018) 161–165.

[12] C. Li, X. Dong, Q. Yuan, G. Xu, Z. Di, Y. Yang, J. Hou, L. Zheng, W. Chen, G. Wu, Identification of novel characteristic biomarkers and immune infiltration profile for the anaplastic thyroid cancer via machine learning algorithms, J Endocrinol Invest, 46 (2023) 1633–1650.

[13] J. Goveia, K. Rohlenova, F. Taverna, L. Treps, L.C. Conradi, A. Pircher, V. Geldhof, L. de Rooij, J. Kalucka, L. Sokol, M. García-Caballero, Y. Zheng, J. Qian, L.A. Teuwen, S. Khan, B. Boeckx, E. Wauters, H. Decaluwé, P. De Leyn, J. Vansteenkiste, B. Weynand, X. Sagaert, E. Verbeken, A. Wolthuis, B. Topal, W. Everaerts, H. Bohnenberger, A. Emmert, D. Panovska, F. De Smet, F.J.T. Staal, R.J. McLaughlin, F. Impens, V. Lagani, S. Vinckier, M. Mazzone, L. Schoonjans, M. Dewerchin, G. Eelen, T.K. Karakach, H. Yang, J. Wang, L. Bolund, L. Lin, B. Thienpont, X. Li, D. Lambrechts, Y. Luo, P. Carmeliet, An Integrated Gene Expression Landscape Profiling Approach to Identify Lung Tumor Endothelial Cell Heterogeneity and Angiogenic Candidates, Cancer Cell, 37 (2020) 21-36.e13.

[14] H. Medfai, A. Khalil, A. Rousseau, V. Nuyens, M. Paumann-Page, B. Sevcnikar, P.G. Furtmuller, C. Obinger, N. Moguilevsky, O. Peulen, M. Herfs, V. Castronovo, M. Amri, P. Van Antwerpen, L. Vanhamme, K. Zouaoui Boudjeltia, Human peroxidasin 1 promotes angiogenesis through ERK1/2, Akt, and FAK pathways, Cardiovasc Res, 115 (2019) 463–475.

[15] V. Castronovo, D. Waltregny, P. Kischel, C. Roesli, G. Elia, J.N. Rybak, D. Neri, A chemical proteomics approach for the identification of accessible antigens expressed in human kidney cancer, Mol Cell Proteomics, 5 (2006) 2083–2091.

